# A Novel Uropathogenic *Escherichia Coli* Genome (strain D3) and Comparative Analysis with Other Uropathogenic and Nonpathogenic Strains

**DOI:** 10.1101/197533

**Authors:** Brian B. Nadel, Shawn J. Cokus, Marco Morselli, Laura J. Marinelli, David Lopez, Robert L. Modlin, Joseph Distefano, Matteo Pellegrini

**Affiliations:** Molecular Cellular and Developmental Biology, University of California, Los Angeles; Dermatology, University of California, Los Angeles; Computer Science, University of California, Los Angeles; Institute of Genomics and Proteomics, University of California, Los Angeles

**Keywords:** Uropathogenic E. Coli, Fimbria, Urinary Tract Infection, Escherichia Coli

## Abstract

**Background:** Bacterial urinary tract infections are extremely prevalent, with half of women having at least one infection at some point in their lives. Most often the causative pathogen is the common gut microbe *Escherichia coli*. One such *E. coli*, strain D3, caused a bladder infection in a male adult, and was resistant to multiple antibiotics. We sequenced and assembled the genome of D3, and present it along with a comparative analysis against other pathogenic and nonpathogenic *E. Coli* strains.

**Results:** By comparing the predicted proteins of D3 with those from 5 uropathogenic and 7 nonpathogenic *E. Coli* strains, we generated a list of 38 genes present in most (4-5) pathogenic strains, but absent in all nonpathogenic strains. Among these were 9 proteins of the Pap fimbrial operon, which has previously been associated with cell adherence and the formation of biofilms. Lastly, we analyzed the list of predicted genes uniquely present in D3 compared to all other strains, and identified multiple transposable elements.

**Conclusions:** The presence of fimbria in most pathogenic *E. coli* strains, and their absence in nonpathogenic ones, suggests that they play a role in pathogenicity, a notion supported by previous work. We also found that D3-specific genes are strongly enriched with transposases, recombinases, and integrase, suggesting that these mobile elements have been inserted or expanded in D3, relative to other strains in the study.

## Background

Urinary tract infections occur frequently even in developed countries, and can result in over 2 million emergency room visits a year in the U.S. alone [1]. Most commonly, the pathogen is a formerly intestinal commensal *E. Coli* that enters the urinary tract externally through the urethra [2]. These uropathogenic *E. Coli* (UPEC) can then travel up the urinary tract into the bladder, ureter, and even kidneys, sometimes resulting in serious medical complications [2]. Thus, understanding UPECs, and the genes that contribute to their pathogenicity, is of direct medical relevance.

In this study, we sequenced and annotated the genome of a UPEC from an adult male patient with a severe, drug resistant infection. In order to identify genetic features unique to this strain, as well as those associated with pathogenicity, we compared its genes to those of several pathogenic and nonpathogenic *E. Coli*. We obtained the genomes of 5 previously sequenced pathogenic and 7 previously sequenced non-pathogenic strains, and determined which proteins are most specific to uropathogenic strains, and thus potentially related to pathogenicity. We found the Pap fimbrial operon to be strongly associated with pathogenic strains.

While a number of adaptations are known to contribute to UPEC’ s survival and virulence, previous work suggests that fimbria may be the most important [3]. Fimbria are extracellular appendages, distinct from flagella or cilia, which allow the bacterium to latch onto host cells or other bacteria [4]. In some cases, fimbria can also allow the bacterium to invade host cells, where it becomes more difficult for the host immune system to detect [3]. Our study supports these previous results and further demonstrate the association of fimbriae with pathogenic strains.

## Results and Discussion

### Genome sequencing, assembly and annotation

D3 is a virulent *E. Coli* strain that caused urinary tract infections in both a 37 year old male and his infant son, and was resistant to the antibiotics nitrofurantoin, sulfamethoxazole, and trimethoprim. In order to explore the basis for D3’ s virulence and antibiotic resistance, we isolated D3 from the urine of the 37 year old male, and sequenced and assembled its genome into 10 contigs representing the main circular genome (4,985,987 bp total) a large plasmid (size 121,318 bp) and a very small, but also very abundant plasmid (1546 bp). Details about assembly are included in the supplementary materials. We ran the FgenesB gene predictor on all D3 contigs and plasmids, and predicted 4873 genes.

The main circular genome of D3 was highly similar to a previously sequenced uropathogenic E. Coli, strain D-i2 [5]. The ten contigs of the main D3 genome aligned well with the D-i2 genome (approximately 99.7% nucleotide similarity), and the D-i2 genome was used to scaffold the contigs of D3. However, since there were over 200,000 bp in D3 that did not align with D-i2, we considered D3 to be a novel strain (Fig 1).

**Figure 1.**
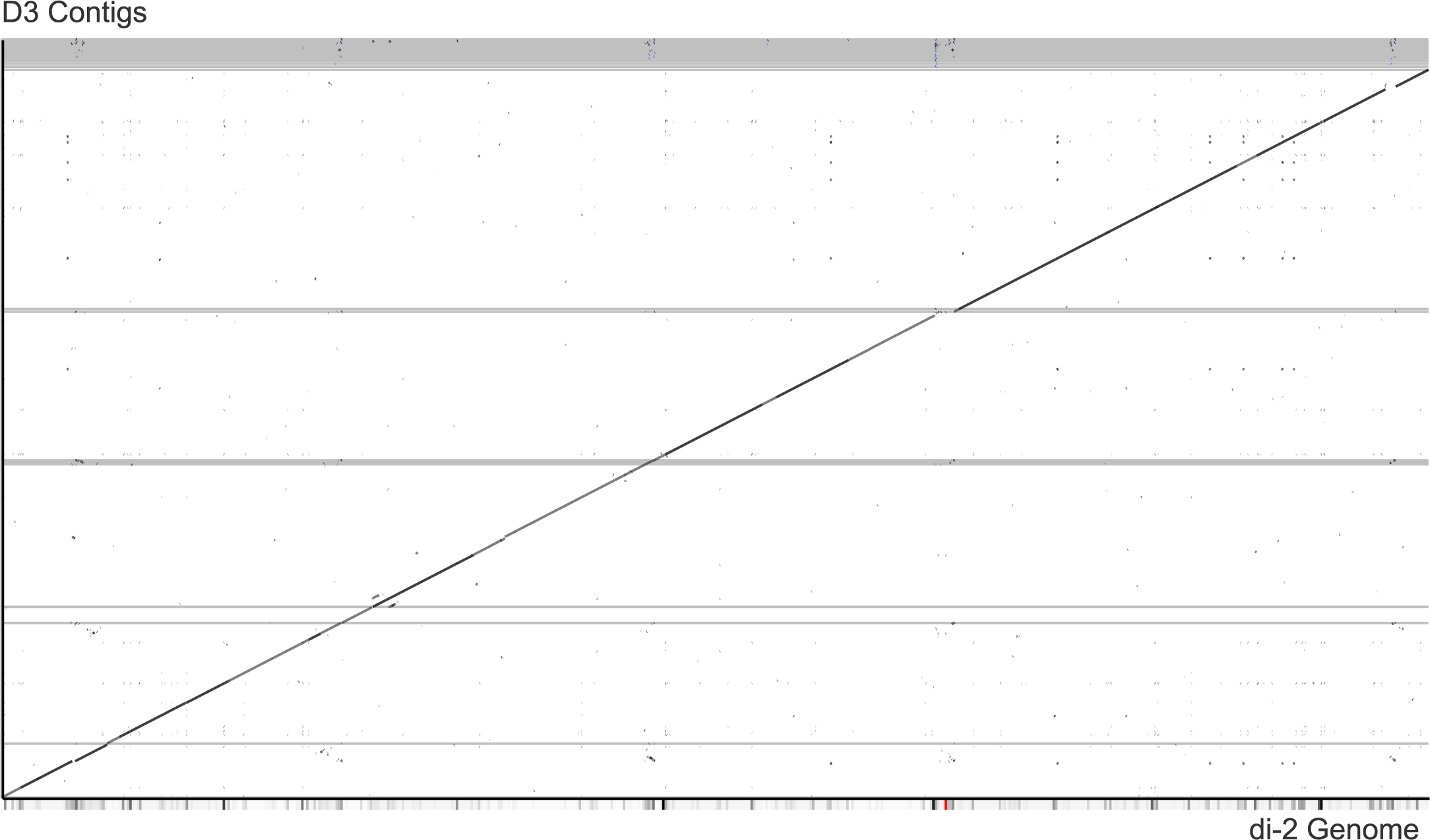
Synteny plot between D3 and D-i2. Horizontal lines represent breaks between D3 contigs. The gray portion at the top of the graph is the result of many small contigs that did not align well with D-i2. Image generated using r2cat [9].

We retrieved a list of 5049 predicted and annotated D-i2 proteins, rRNAs and tRNAs from NCBI. D3 contained exact matches (at the protein level for genes, nucleotide for RNAs) for 4566 of these, and 4799 were more than 98.5% similar. After aligning the D-i2 genome with the 10 D3 contigs, we transferred annotations for these 4799 genes/RNAs that were >98.5% identical from the D-i2 predicted protein (and RNA) list.

### Putative Pathogenicity genes

We obtained the predicted protein lists of 12 *E. Coli* strains from NCBI, and compared these lists with the predicted protein list of D3 (i.e. the output of FGenesB) in order to generate a list of putative pathogenic proteins. 5 of the protein lists obtained from NCBI were from nonpathogenic E. coli strains, While the remaining 7 were from uropathogenic strains (criteria for selecting these genomes is described in methods).

We considered 2 genes to be orthologous if they passed the mutual best hit orthology test using the bit score of blastp. Using this definition, we computed phylogenetic profiles that describe the occurrence of orthologs for each protein in D3 across the other 12 strains (Table 1 describes the number of orthologs present in each strain). We then isolated a set of 38 genes in D3 that have orthologs and inat least 4 other uropathogenic strains, but no nonpathogenic strains (see additional file 1 for orthology spreadsheet). We then annotated this list by BLASTing against the NCBI nr (see additional file 2 for gene list with annotations). To provide a visual representation of the functional distribution of these proteins, we also ran the sequences of these genes through Blast2GO (Fig 2a), and manually categorized them by keyword search (Figure 2b) [7].

**Table 1.** Number and percentage of predicted genes in D3 that had orthologs in other strains. There were 4873 predicted genes total, and presence of orthologs was determined by mutual best hit orthology tests.

| Strain | D3 Genes with Orthologs | Percentage with Orthologs |
| --- | --- | --- |
| D3 | 4873 | 100.00% |
| Di2 | 4363 | 89.53% |
| CFT073 | 4321 | 88.67% |
| IAI39 | 3767 | 77.30% |
| NA114 | 3975 | 81.57% |
| UTI189 | 4123 | 84.61% |
| DH1 | 3625 | 74.39% |
| DH10B | 3500 | 71.82% |
| ED1a | 3875 | 79.52% |
| MG1665 | 3618 | 74.25% |
| REL606 | 3597 | 73.81% |
| SE15 | 3952 | 81.10% |
| W3110 | 3630 | 74.49% |

**Figure 2.**
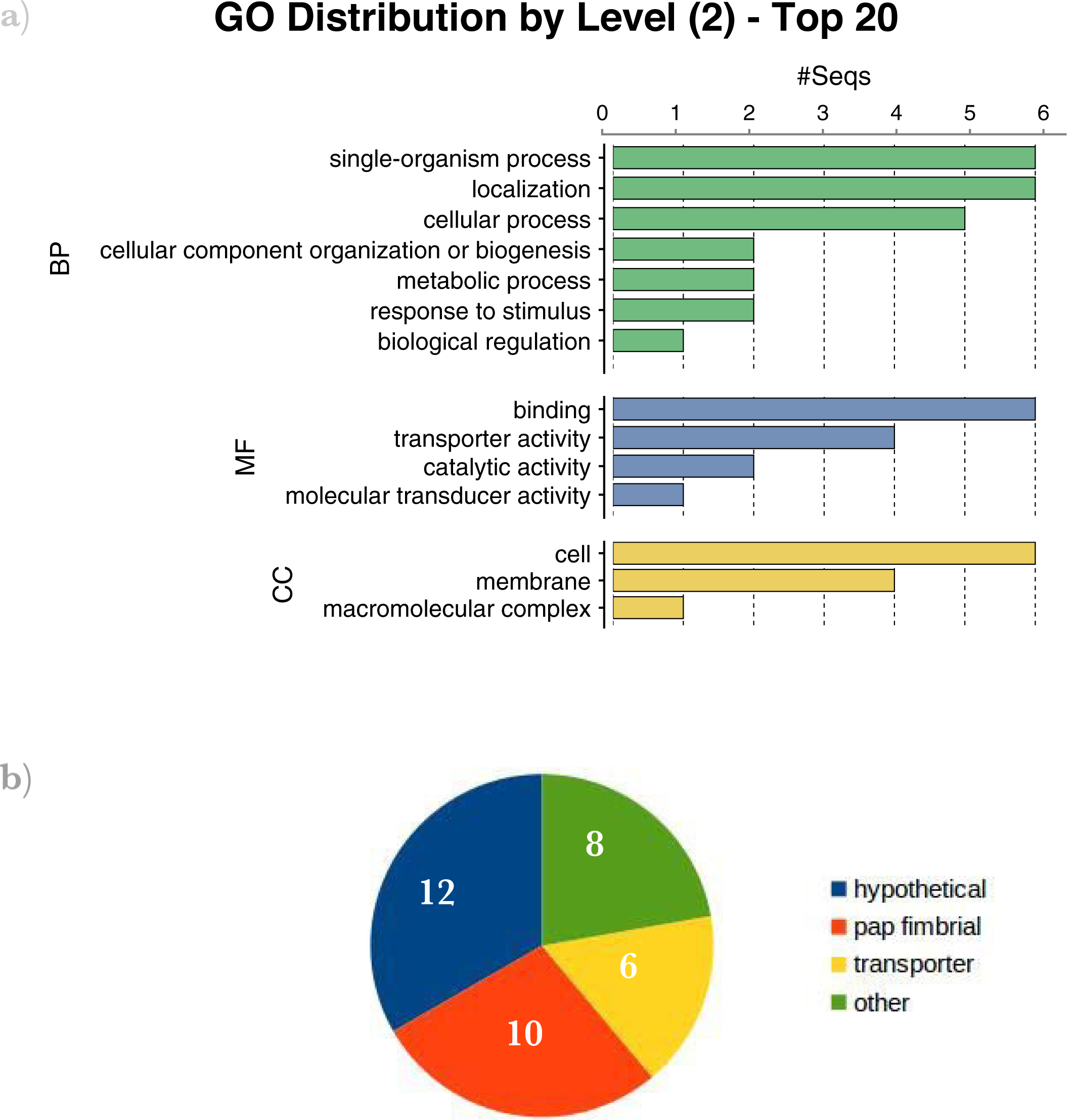
a) Functional categories of genes present in 4-5 pathogenic strains, and 0 nonpathogenic strains. Graph produced by Blast2Go with default parameters [7]. BP = Biological Process, MF = Molecular Function, CC = Cellular Component. b) keyword-based categorization of annotations.

This list contained 10 Pap fimbrial genes,, which code for fimbrial proteins known to bind to the mammalian kidney [8]. An 11th gene, PapJ was also present in D3, and had orthologs in 3 pathogenic strains, and 0 nonpathogenic strains. In D3, 10 of these genes appear to lie in an operon; they are found within a stretch of 9070 base pairs, and no other predicted genes are found in that stretch (see figure 3). The 11^th^ gene, PapB lies on a seperate contig than the other Pap genes. A STRING [9] analysis of the Pap genes reveals that the Pap genes are present in close proximity (within 300 bp) in many species of bacteria (see additional file 3).

**Figure 3.**
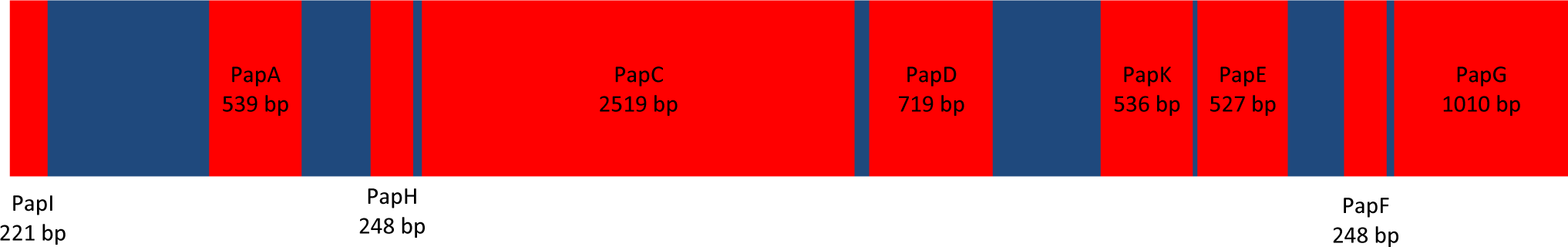
The Pap operon as it appears in D3. All nine sequences in D3 were >94% identical to the corresponding Pap genes, and all except PapA were >98% identical. PapI lies on opposite strand from other genes in operon.

D-i2 is the only pathogenic strain in the study that does not contain orthologs for the 11 pap fimbrial genes found in D3 (with the exception of PapB, which does have an ortholog). This is unexpected, due to the high sequence similarity between the two strains. However, a number of other fimbrial genes were present in D-i2, including 9 consecutive genes with roles related to F1C fimbria. It is possible that this F1C operon fulfills a role similar to that of the pap operon.

This list also contained a 3 gene phosphotransferase system operon (see figure 4). It is unclear whether this system has a role in pathogenicity. Membrane proteins were also common in this list, as were hypothetical proteins.

**Figure 4.**
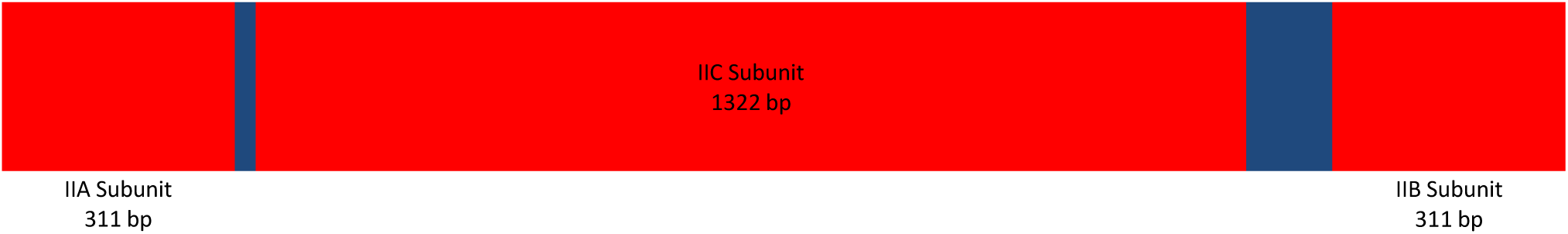
The lactose/cellobiose phosphotransferase operon as it appears in D3. All 3 sequences in D3 were > 99% identical to best BLAST hits.

### D3 Specific genes

We next sought to identify genes that were unique to the D3 strain, compared to the other 12 *E. coli* strains. We identified 115 predicted genes in D3 that did not have orthologs in any of the 12 other *E. Coli* genomes in the study. We annotated these genes by BLASTing against the NCBI nr database (additional file 4). These predicted genes had varied functions ( figure 5). 15 of these predicted genes were described as integrases, recombinases, or transposases. There were also 15 proteins described as transporters, 5 conjugal transfer proteins, and 4 plasmid-related proteins. 25 of these predicted genes had no good BLAST hits to the nr protein database, or only hits to unannotated genes. These 25 predicted genes may be pseudogenes, or perhaps have not yet been described and deposited to the NCBI database.

**Figure 5.**
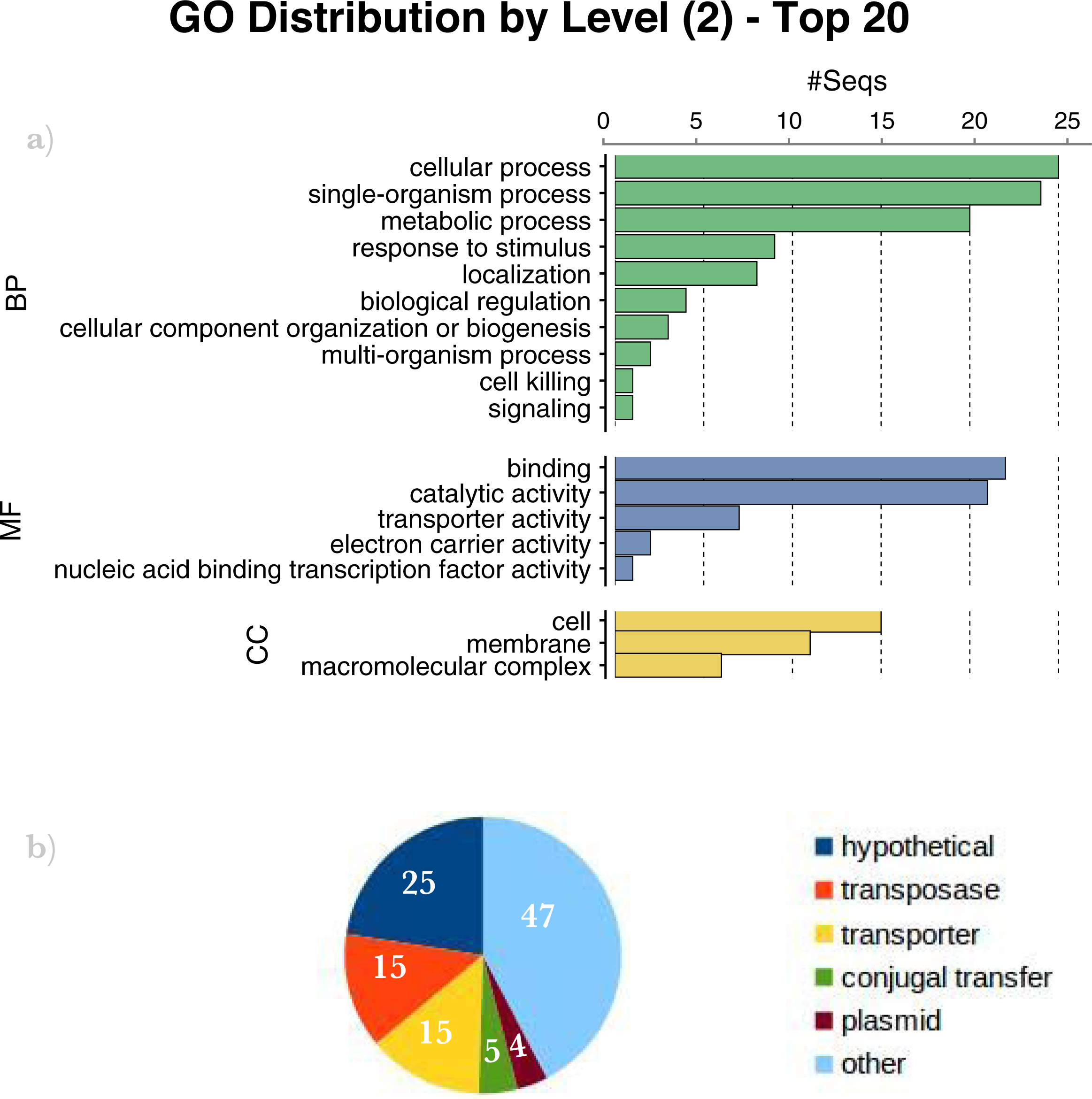
a) Functional categories of genes present in D3, but no other strains in the study. Graph produced by Blast2Go with default parameters [7]. BP = Biological Process, MF = Molecular Function, CC= Cellular Component. b) keyword-based categorization of annotations.

### Conclusions

We sequenced a UPEC strain we obtained from a drug resistant infection in an adult male. After assembling the genome and running gene prediction software, we found that the strain was very similar to a previously sequenced strain, D-i2. However, we identified 115 genes that were present in D3, but absent in D-i2, as well as the genomes of the 11 other *E. Coli* strains used in the study. When comparing this D3-specific gene list to the large portion of the genome annotated by alignment with D-i2, we found that the D3 specific genes were heavily enriched with transposases, recombinases, and integrases. In the entire D3 genome, there were 29 of these genes, while the D3-specific list contained 17. This results in a rate of.4% in the total genome, and 16.5% in the D3 specific list. This suggests that these mobile elements have been inserted, and possibly expanded, in D3 compared to D-i2.

Our results also suggest that fimbriae may be an important adaptation for uropathogenicity, a notion supported by previous work [3]. In the present study, 4 of the 5 pathogenic strains contained orthologs to the D3 pap fimbrial operon, and the 5th strain contained a (non-pap) fimbrial operon, as well. The distribution of these pap genes (i.e. their prevalence among pathogenic strains and absence in nonpathogenic strains) is suggestive of a pathogenic function, though not conclusive due to the associative nature of the study. It is interesting that D-i2 does not contain an ortholog for the D3 pap operon, as 92% of its genes exactly match genes found in D3, and many of the remaining genes are very similar. This could mean that D3 and D-i2 acquired pathogenicity independently of one another, though further work would need to be done to test this hypothesis.

## Methods

### Sequencing and Assembly

D3 was isolated from the urine of a 37 year-old male who was diagnosed with a urinary tract infection. It was cultured on Luria-Bertani (LB) agar or in liquid LB medium at 37°C. Bacterial DNA was prepared from an overnight culture using the Wizard→?Genomic DNA Purification Kit (Promega), according to manufacturer’s instructions for the isolation of Genomic DNA from Gram-negative bacteria.

DNA of two different extraction methods was fragmented using Covaris. Illumina TruSeq DNA kit was used to prepare the library following manufacturer instructions. Briefly, fragmented DNA was end-repaired and A-tailed. Differently barcoded TruSeq adapters were ligated to each sample followed by PCR amplification. Samples were sequenced with 100 paired-end strategy using the Illumina HiSeq2000 platform. The libraries were then assembled using ABySS. A detailed description of the assembly is presented in the supplementary materials.

### Obtaining Gene Lists

We predicted a set of 4873 likely genes using the FgenesB program of the Molquest package (version 2.4.5.1135), which we obtained from the Softberry website (http://linux1.softberry.com/berry.phtml). We ran FgenesB on all D3 contigs with unmodified parameters, other than the required options (organism: Escherichia coli K-12, translation table: Bacterial and Plant Plastid, Trash: 60=default).

In addition to D3, we used protein lists from 12 fully sequenced E. Coli strains obtained from the NCBI database (see table 2 for accession information). Five of these strains were uropathogenic, which included D-i2, CFT073, UTI189, NA114, and IAI39. These strains were explicitly described by NCBI as uropathogenic, or as isolated from a bladder infection. We used all strains that were described as such and available at the time of the study, and which had complete genomes. The 7 nonpathogenic strains were DH1, DH10B, ED1a, MG1665, REL606, SE15, and W3110 (figure 4). 4 of these were lab strains that were not described as pathogenic, while the rest were described as either “nonpathogenic” or “avirulent”.

**Table 2.**
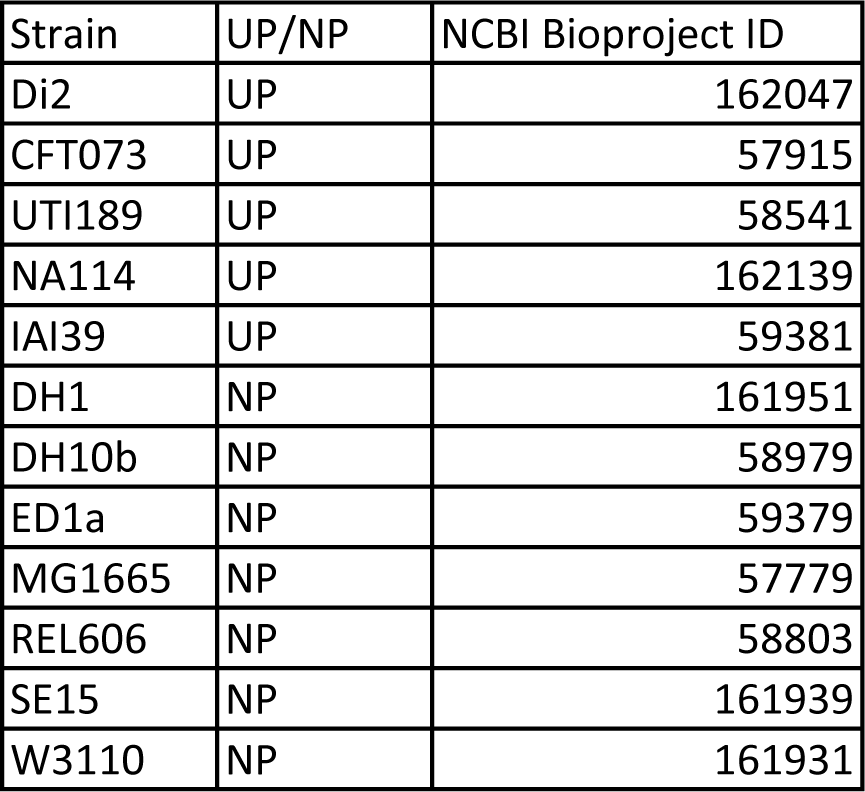
Accession information for strains used in the study. UP = Uropathogenic, NP = Nonpathogenic.

### Annotation and Comparative Analysis

Given the protein list from D-i2 (which was previously annotated) we transferred 4799 annotations from D-i2 to D3. Within the 10 contigs that aligned well with the D-i2 genome, we transferred annotations for all genes that had >98.5% identical amino acid sequences.

We identified putative pathogenicity genes, which we defined as genes that were present in D3 and had orthologs in 4 (or more) of the 5 pathogenic strains, and none of the nonpathogenic strains. Our criteria for determining whether a D3 gene had an ortholog in another strain was mutual best hit of the bit score of blastp. We then manually annotated the resulting 41 genes by BLASTing against the NCBI nr database. We chose the most specific annotation among the best few hits, so long as sequence identity and coverage were both greater than 96% (one or both were 100% in most cases).

In the same manner, we also annotated predicted genes that were present in D3, but not in any other strains in the study. As this list was relatively long (600 predicted genes), we excluded any proteins less than 100 amino acids in length. 127 predicted proteins remained.

## Availability of Supporting Data

Supporting data is available through NCBI (BioProject ID: PRJNA252526; BioSample accession number: SAMN02851119; Sequence Accession number pending). Additional file 1 contains an orthology spreadsheet describing the presence of orthologs between D3 and the other strains in the study. Additional file 2 is a list of putative pathogenic genes, including their annotations. Additional file 3 contains the results of a STRING analysis on the Pap fimbrial genes. Additional file 4 is an annotated list of D3 predicted genes that did not have orthologs in any of the other strains in the study.

## List of Abbreviations

*E. Coli*: Escherichia Coli
UPEC: Uropathogenic *Escherichia Coli*

## Competing Interests

The authors declare that they have no competing interests.

## Author’s Contributions

BN analyzed the assembled genome, and drafted the manuscript. SC sequenced and assembled the genome, and assisted with the post-assembly analysis. RM and LJM isolated and grew the *E. Coli* samples and extracted their DNA. MM prepared the sequencing libraries for the samples. JD provided the samples and initiated the study. DL deposited the sequence data and helped make the figures. MP supervised the project, and helped draft the manuscript.

## Acknowledgements

We received funding from the P50 AR063020-01 grant.

